# RNA degradation sculpts the maternal transcriptome during *Drosophila* oogenesis

**DOI:** 10.1101/2020.06.30.179986

**Authors:** Patrick Blatt, Siu Wah Wong-Deyrup, Alicia McCarthy, Shane Breznak, Matthew D. Hurton, Maitreyi Upadhyay, Benjamin Bennink, Justin Camacho, Miler T. Lee, Prashanth Rangan

## Abstract

In sexually reproducing animals, the oocyte contributes a large supply of RNAs that are essential to launch development upon fertilization. The mechanisms that regulate the composition of the maternal RNA contribution during oogenesis are unclear. Here, we show that a subset of RNAs expressed during the early stages of oogenesis is subjected to regulated degradation during oocyte specification. Failure to remove these RNAs results in oocyte dysfunction and death. We identify the RNA-degrading Super Killer complex and No-Go Decay factor Pelota as key regulators of oogenesis via targeted clearance of RNAs expressed in germline stem cells. These regulators target RNAs enriched for cytidine sequences bound by the protein Half pint. Thus, RNA degradation helps orchestrate a germ cell-to-maternal transition by sculpting the maternal RNA contribution to the zygote.

## Report

A fertilized egg is totipotent, having the unique potential to differentiate into every cell lineage in the adult organism^1-3^. Across animals, 40-75% of genes are deposited into the egg during oogenesis as part of the maternal RNA contribution required for embryo development^4-6^. It is unlikely that every RNA synthesized during oogenesis is destined for the maternal contribution: RNAs that support oogenesis-specific functions, such as germline stem cell (GSC) self-renewal and differentiation, could be detrimental during embryogenesis. It is not known if such oogenesis-specific RNAs are targeted for elimination or what, if any, mechanisms ensure that only the appropriate RNAs are deposited into the oocyte.

In *Drosophila*, oogenesis occurs in ovarioles composed of germaria, which contain the GSCs and the GSC daughter cells (cystoblasts, CB) that progressively differentiate into 16-cell cysts (Figure 1A)^7-11^. In each cyst, the oocyte receives RNA and protein contributions from the remaining 15 nurse cells (Figure 1A’), thus causing the oocyte to enlarge forming egg chambers (Figure 1A)^12-18^. In a screen to identify novel regulators of this process, we discovered that a component of the RNA-degradation-promoting Super Killer (Ski) complex (Figure 1B), Super Killer 2 (Ski2), called Twister (Tst) in *Drosophila*, is required for egg chamber growth and female fertility (Figure S1A)^19-22^. Wild type (WT) *Drosophila* ovarioles stained for Vasa (germ cells) and for 1B1 (somatic cell membranes) show the progression from the germarium to successively larger egg chambers (Figure 1C). In contrast, egg chambers failed to grow in *tst* mutant ovarioles (Figure 1C-D, 1M) as well as upon germline RNAi depletion of *tst* (*nanos-GAL4 >*RNAi, Figure 1E-F, 1M) but not when *tst* was depleted in the soma (*traffic jam-GAL4 >*RNAi, S1B-C)^23,24^. However, *tst* mutant flies are otherwise viable, and successful oogenesis and egg production were restored in *tst* mutants by expressing Tst protein in the germline alone (Figure 1G-H, 1M, S1A). Egg chambers lacking *tst* expressed cleaved Caspase 3 at putative stages 6-7, suggesting that they undergo apoptosis (Figure S1D-E).

**Figure 1.**
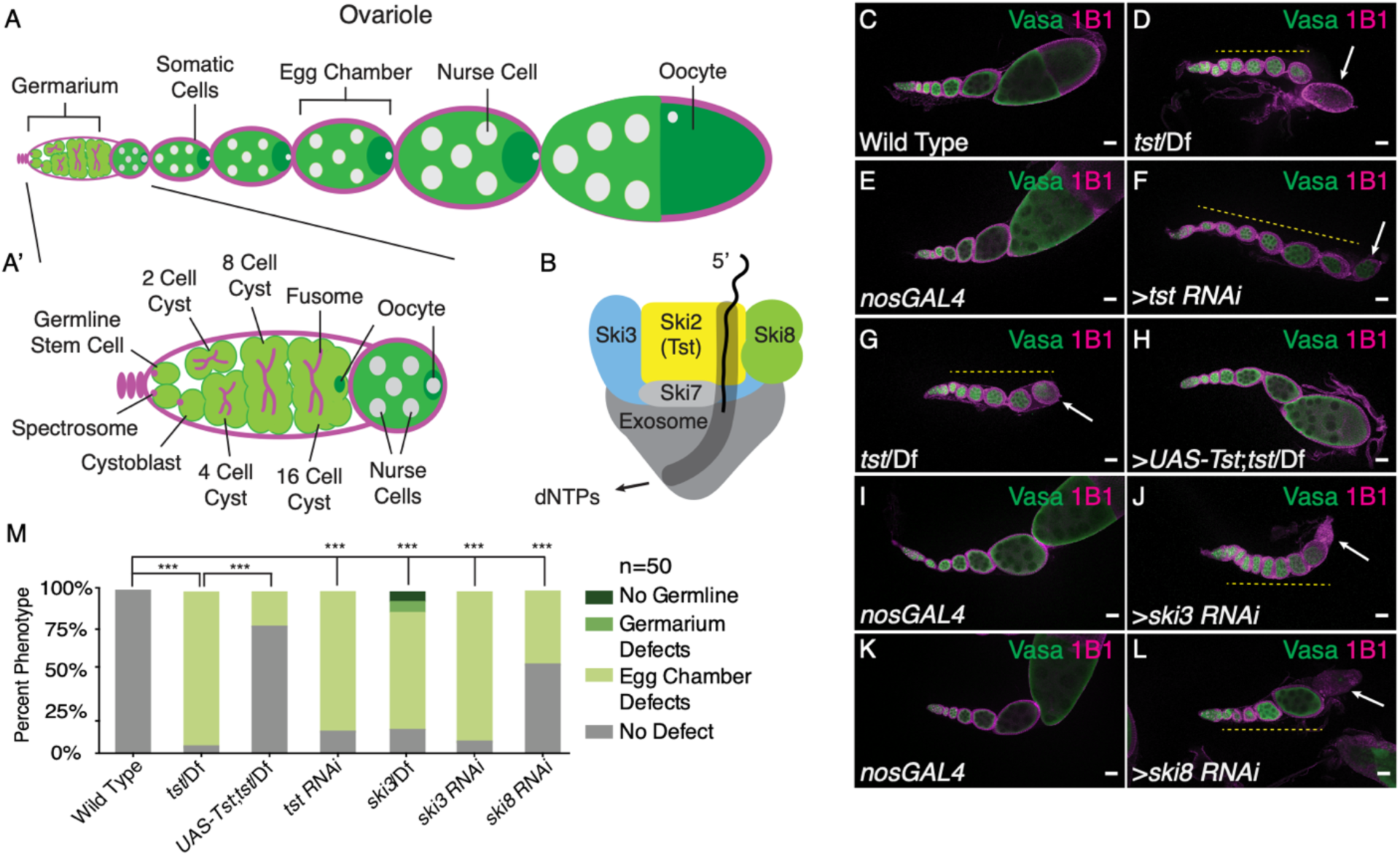
Components of the Ski Complex are required in the germ line for oogenesis. (**A**) Schematic of a *Drosophila* ovariole and (**A’**) germarium. (**B**) Representation of the Ski Complex composed of Ski2 (Tst, yellow), Ski3 (blue), two Ski8 proteins (green), and Ski7 (light gray) threading mRNA (black) into the exosome (dark gray) where mRNA degradation occurs. (**C**) Confocal image of an adult WT control ovariole stained with Vasa (green) and 1B1 (magenta) showing normal egg chamber development. (**D**) Confocal image of a *tst* genomic mutant ovariole stained with 1B1, Vasa and indicating egg chambers that do not grow in size (yellow dashed line) and dying egg chamber (arrow). (**E**) Confocal image of a *nosGAL4* driver control ovariole stained with 1B1 and Vasa. (**F**) *tst* germline RNAi knockdown ovariole stained with 1B1, Vasa, and indicating egg chambers that do not grow in size (yellow dashed line) and dying egg chamber (arrow). (**G**) Confocal image of a *tst* genomic mutant ovariole stained with 1B1 and Vasa. (**H**) Confocal image of a *tst* genomic mutant ovariole expressing recombinant Tst protein in the germline stained with 1B1 and Vasa. (**I**) Confocal image of a *nosGAL4* driver control ovariole stained with 1B1 and Vasa. (**J**) *ski3* germline RNAi knockdown ovariole of stained with 1B1, Vasa, and indicating egg chambers that do not grow in size (yellow dashed line) and dying egg chamber (arrow). (**K**) Confocal image of a *nosGAL4* driver control ovariole stained with 1B1 and Vasa. (**L**) *ski8* germline RNAi knockdown ovariole stained with 1B1, Vasa, and indicating egg chambers that do not grow in size (yellow dashed line) and dying egg chamber (arrow). (**M**) Quantification of oogenesis defect phenotypes observed in Ski complex genomic mutants, germline RNAi knockdowns and UAS-Tst rescue (Control vs *tst*/Df n=50, p<0.001, *tst*/Df vs *UAS-Tst*;*tst*/Df n=50, p<0.001, Control vs *tst RNAi* n=50, p<0.001, Control vs *ski3*/Df n=50, p<0.001, Control vs *ski3 RNAi* n=50, p<0.001, Control vs *ski8 RNAi* n=50, p<0.001, Chi Square Analyses). Scale bars are 10μm.

The Ski complex, in addition to RNA helicase SKI2, consists of the scaffolding subunits SKI3 and SKI8, which are coupled to the exosome complex by SKI7 (Figure 1B)^20,21,25,26^. We found that *ski3* (*CG8777*) mutant and germline depletion of *ski3* and *ski8* (*CG3909*) phenocopied *tst* mutants (Figure 1I-M, S1F-H). HBS1 is thought to fulfill the role of SKI7 in *Drosophila*; however, female *hbs1* mutants were previously found to be fertile, suggesting that SKI7/HBS1 is dispensable for Ski complex function in the female germline or acts redundantly with a yet-unidentified protein^27-29^. Overall, we conclude that the Ski complex components Ski2, Ski3 and Ski8 are required in the fly germ line for oogenesis.

Given the role of the Ski complex in exosome-mediated RNA degradation, we hypothesized that Tst promotes degradation of RNAs during oogenesis^21,30,31^. RNA sequencing (RNA-seq) revealed 296 genes upregulated in ovaries lacking *tst* (Figure 2A, Supplemental Table 1). These include 207 genes such as *blanks* and actin 57B (*act57B*) with >4-fold higher levels in a germline *tst RNAi* compared to WT (Figure 2B), which likely represent transcripts regulated by Tst in the germline. To determine that the depletion of *tst* resulted in a defect in post-transcriptional regulation, we measured pre-mRNA levels of select Tst-regulated RNAs by qRT-PCR and indeed found no significant difference between WT and *tst* germline RNAi flies (Figure S2A). Taken together, these data suggest that *tst* promotes the post-transcriptional degradation of a distinct group of RNAs during oogenesis.

**Figure 2.**
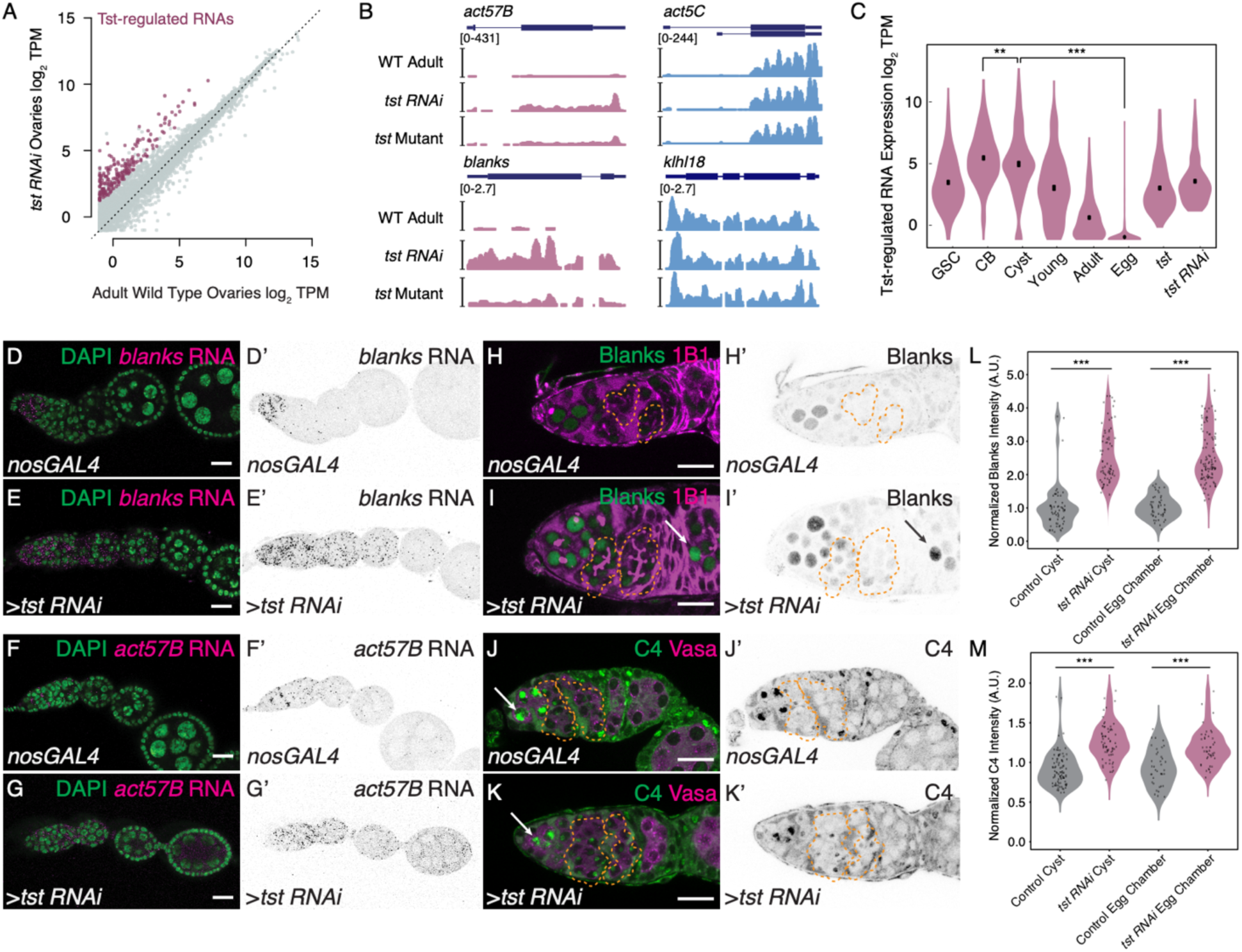
Tst promotes degradation of a subset of transcripts prior to oocyte specification. (**A**) Biplot of RNA-Seq data from adult WT and *tst* germline RNAi knockdown ovaries in log_2_ Transcripts Per Million (TPM) highlighting upregulated Tst-regulated RNAs (magenta). (**B**) Genome browser tracks of Tst-regulated genes *act57B* and *blanks* (magenta) and non-target genes *act5C* and *klhl18* (blue). (**C**) Violin plot of Tst-regulated RNAs from germline stem cells (GSC), cystoblasts (CB), differentiating cysts (Cyst), young WT ovaries (Young), adult WT ovaries (Adult), unfertilized eggs (Egg), *tst* mutant (*tst*) and *tst* germline RNAi (*tst RNAi*) ovaries showing the decrease in expression of Tst-regulated RNAs after differentiation and cyst stages (n=207, CB vs cyst p<0.002, cyst vs egg p<0.0001, Paired t-Test). (**D-D’**) Confocal images of *in situ* hybridizations probing against *blanks* RNA (magenta, grayscale) and DAPI (green) in *nosGAL4* showing *blanks* RNA expression restricted to the undifferentiated cells and in (**E-E’**) *tst RNAi* ovarioles where *blanks* RNA expression is expanded to egg chambers. (**F-F’**) Confocal images of *in situ* hybridizations probing against *act57B* mRNA (magenta, grayscale) and DAPI (green) in *nosGAL4* showing low *act57B* RNA expression and (**G-G’**) *tst RNAi* ovarioles exhibiting expanded *act57B* RNA expression in the germarium and egg chambers. (**H-H’**) Confocal images of *nosGAL4* and (**I-I’**) *tst RNAi* germaria stained for 1B1 (magenta) and Blanks protein (green and grayscale) showing expanded Blanks expression in *tst RNAi* cysts (orange dashed lines) and egg chambers (arrow). (**J-J’**) Confocal images of *nosGAL4* and (**K-K’**) *tst RNAi* germaria stained for Vasa (magenta) and C4 antibody (nuclear Actin) (green and grayscale) showing expanded nuclear Actin expression in *tst RNAi* cysts (orange dashed line). (**L**) Arbitrary Units (A.U.) quantification of Blanks protein expression normalized to control cysts in Control (gray) and *tst RNAi* (magenta) cysts and egg chambers (WT cyst n=60, *tst RNAi* cyst n=70, p<0.0001. WT egg chamber n=49, *tst RNAi* egg chamber n=102, p<0.0001, Student’s t-Test). (**M**) A.U. quantification of nuclear Actin expression normalized to control cysts in Control (gray) and *tst RNAi* (magenta) cysts and egg chambers (WT cyst n=92, *tst RNAi* cyst n=73, p<0.0001. WT egg chamber n=38, *tst RNAi* egg chamber n=45, p<0.0001, Student’s t-Test). Scale bars are 10μm.

To determine when Tst acts, we used RNA-seq to profile the expression of Tst-regulated RNAs in ovaries across a time course of oocyte development: GSCs, CBs, and cysts, which were each enriched using mutants (see Methods); germaria and early egg chambers were enriched using young WT ovaries; late-stage egg chambers were enriched using adult WT ovaries; and unfertilized eggs, which represent the maternal contribution^9,11,32-34^. Principal component analysis revealed that *tst* mutant and *tst RNAi* ovaries more closely resemble WT as compared to undifferentiated stages, suggesting that *tst* is required after differentiation (Figure S2B). Indeed, compared to non-targets, Tst-regulated RNAs decreased at the cyst stages and were nearly absent as part of the maternal contribution in the egg (Figure 2C, S2C-D). *In situ* hybridization of the Tst-regulated RNAs *blanks* and *act57B* demonstrates low levels beginning in the cyst stages in WT, in contrast to persistence throughout the egg chambers in *tst* germline RNAi (Figure 2D-G’). To precisely determine when Tst-regulated RNAs are degraded, we probed for proteins encoded by Tst-regulated RNAs *blanks* and *actins*. In WT, both Blanks and nuclear-Actins (detected by C4 staining) were highly expressed in GSCs and CBs but their expression is attenuated in the cysts, when the oocyte is specified, consistent with previous reports (Figure 2H-M)^35-38^. In contrast, both Blanks and nuclear-Actin expression persisted in the cysts and egg chambers of *tst* germline RNAi flies (Figure 2H-M). We did not find gross changes to cytoplasmic Actin pool upon the loss of *tst*, as measured by Phalloidin staining (Figure S2E-F’)^39^. Overall, our data suggest that Tst attenuates the levels of Blanks and Actin proteins by degrading their mRNAs before oocyte specification.

To investigate how specific transcripts are targeted by Tst, we considered the contribution of RNA surveillance pathways, which are known to direct RNAs to the Ski complex for degradation^40^. Nonsense mediated decay (NMD) and non-stop decay (NSD) are unlikely to be involved. In contrast to *tst, ski3* and *ski8* germline RNAi flies, germline mutant clones of the NMD pathway components *up-frameshift 1* (*Upf1*), *Upf2*, and *Upf3* do produce eggs, albeit with patterning defects^41^. We additionally looked for features in the RNAs that could trigger NMD or NSD. Most Tst-regulated RNAs do not encode introns in their 3’ untranslated regions (3’ UTR) (Figure S3A), nor show any evidence for aberrant splicing that would give rise to premature termination codons (Figure S3B), ruling out NMD^42-44^. NSD is triggered by ribosome read through into the 3’ UTR, but all Tst-regulated RNAs are annotated transcripts that encode stop codons suggesting that NSD is also not involved^45-47^.

However, we did find evidence that no-go decay (NGD), which is activated when ribosomes stall on RNAs, was involved in the degradation of Tst-regulated RNAs. Pelota (Pelo/DOM34) is a critical effector protein of the NGD pathway that promotes recycling of stalled ribosomes on mRNAs^27,48,49^. Intriguingly, *pelo* mutants, like *tst* mutants, are homozygous viable but female sterile, and this role is germline specific^50^. *pelo* mutant egg chambers failed to grow and died mid-oogenesis, phenocopying *tst* mutant ovaries (Figure 3A-C). In addition, *pelo* mutants also lost GSCs, as previously described (Figure S3C-D)^50^. To test if *pelo* and *tst* co*-*regulate target RNAs, we performed RNA-seq on *pelo* mutant ovaries and found that 81% of genes upregulated upon the loss of *tst* were also upregulated >2-fold in *pelo* mutants, including *act57B*, though not *blanks* (168/207, Figure 3D). These data suggest that *pelo*, a key component of the NGD pathway, promotes the degradation of a large fraction of Tst-regulated RNAs.

**Figure 3.**
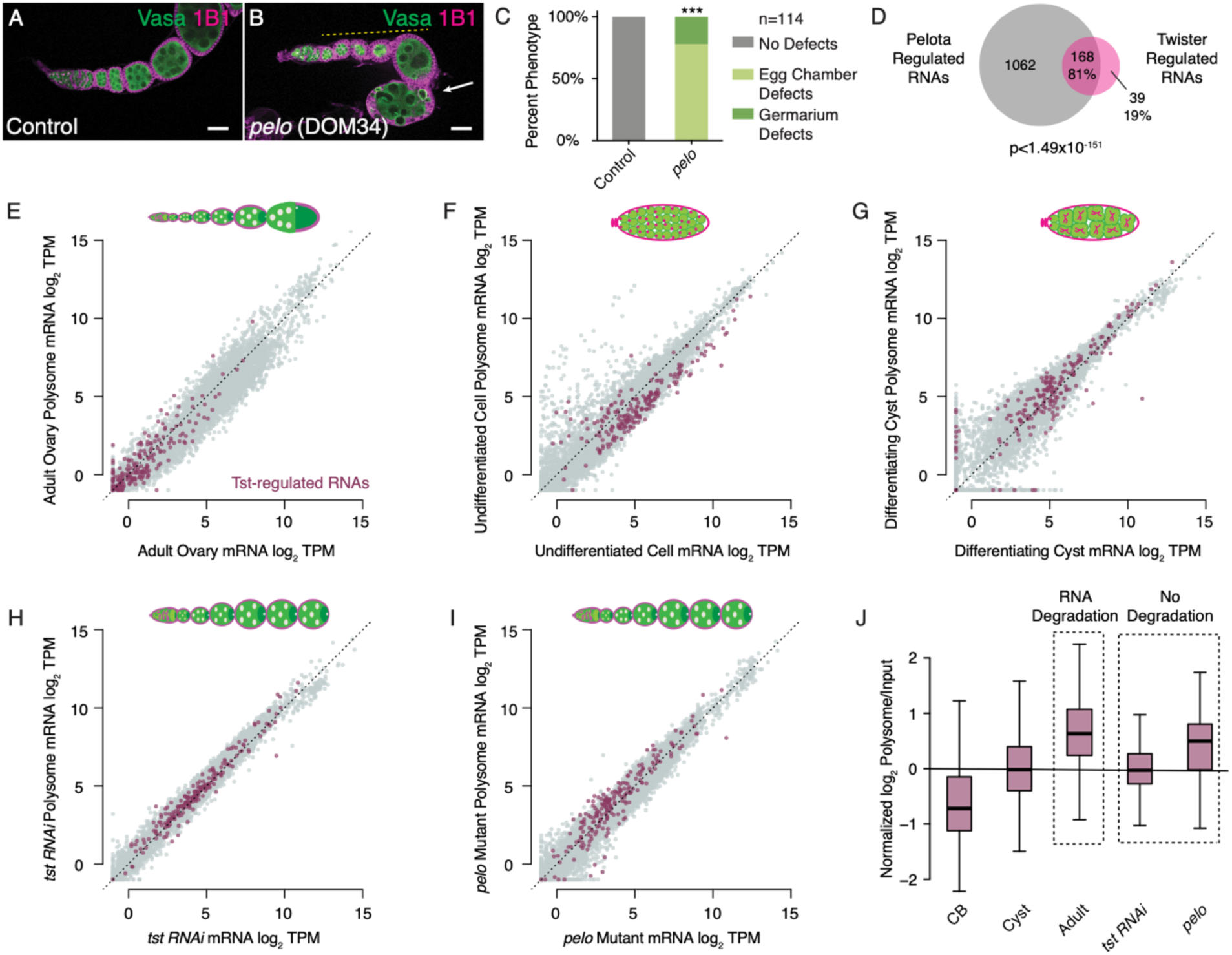
Tst-regulated RNAs are coregulated by Pelota and exhibit an increased ribosome association concurrent with a decrease in mRNA abundance. (**A**) Confocal image of a WT control and (**B**) *pelo*^*1*^ mutant ovariole stained with 1B1 (magenta) and Vasa (green) and indicating egg chambers that fail to grow (yellow dashed line) and subsequently die (arrow). (**C**) Quantification of oogenesis defect phenotypes observed in *pelo*^*1*^ mutants (n=113, p<0.001, Chi Square Analysis). (**D**) Venn diagram illustrating overlap of 81% of Tst-regulated RNAs that are >2 fold upregulated upon loss of *pelo* (p<1.49×10^−151^, Hypergeometric Test). (**E**) Biplot of poly(A) mRNA Input log_2_ TPM versus polysome associated mRNA log_2_ TPM from adult WT ovaries highlighting Tst-regulated RNAs (magenta) showing low RNA abundance. (**F**) Biplot of poly(A) mRNA Input log_2_ TPM versus polysome associated mRNA log_2_ TPM from undifferentiated germ cells highlighting Tst-regulated RNAs (magenta) indicating both an increased RNA abundance and ribosome association compared to Adult WT. (**G**) Biplot of poly(A) mRNA input log_2_ TPM versus polysome associated mRNA log_2_ TPM from differentiating cysts highlighting Tst-regulated RNAs (magenta) indicating both an increased RNA abundance and ribosome association compared to Adult WT. (**H**) Biplot of poly(A) mRNA input log_2_ TPM versus polysome associated mRNA log_2_ TPM in germline *tst RNAi* ovaries highlighting Tst-regulated RNAs (magenta) indicating both an increased RNA abundance and ribosome association compared to Adult WT. (**I**) Biplot of poly(A) mRNA input log_2_ TPM versus polysome associated mRNA log_2_ TPM in *pelo*^1^ ovaries highlighting Tst-regulated RNAs (magenta) indicating both an increased RNA abundance and ribosome association compared to adult WT. Scale bars are 10μm. (**J**) Quantification of normalized log_2_ polysome/input mRNA of Tst-regulated RNAs in CB, cyst, adult, *tst RNAi* and *pelo* samples showing increased ribosome association during the transition from CB to cyst to adult. Ribosome association is comparable for cyst and *tst RNAi* and adult and *pelo* in which RNA degradation is not occurring.

To observe the translation dynamics of Tst-regulated RNAs, we purified polysomes from ovaries enriched for different stages of oocyte development and performed RNA-seq (Figure 3E-G). Adult WT ovaries overall show proportional RNA-seq read depth in the polysome fraction (y-axis) compared to total RNA (x-axis), with Tst-regulated RNAs recapitulating the low expression we observed previously (Figure 3E). In undifferentiated CBs, where Tst-regulated RNA levels are higher, we observed weak polysome association (Figure 3F, J), but overall these RNAs appear to be translated; this is consistent with detection of Blanks protein in CBs prior to oocyte specification (Figure 2H-H’). In the differentiating cysts, polysome association appears to increase (Figure 3G, J); however, we did not observe Blanks protein in WT cysts and egg chambers (Figure 2H-H’), suggesting that ribosome engagement of these Tst-regulated RNAs is not productive. Although increased association with ribosomes is usually linked to increased RNA stability, Tst-regulated RNAs showed an increased association with ribosomes concomitant with their degradation (Figure 3E-G, J)^51^. This change in polysome association is not seen for non-targets (Figure S3E). These results suggest ribosomes are stalled on Tst-regulated RNAs prior to their degradation.

As DOM34 (*pelo*) promotes recycling of stalled ribosomes, and SKI2 is both required to extract RNAs from stalled ribosomes and to promote their degradation, we predicted that Tst-regulated RNAs would be associated with polysomes but not degraded in the later stages of oogenesis in *tst* and *pelo* mutants^52-54^. Indeed, we found that Tst-regulated RNA abundance was substantially increased in *tst* and *pelo* mutant ovaries compared to developmentally similar WT ovaries, and that these RNAs are associated with ribosomes (Figure 3H-J). Taken together, we find that Tst-regulated RNAs have increased association with ribosomes prior to their degradation and are regulated by the NGD pathway member *pelo.*

Pelo-mediated degradation can be activated by features in the coding sequence that cause ribosome stalling, such as sub-optimal codons^55,56^. We found that Tst-regulated RNAs in fact have an elevated codon optimality compared to non-targets, as measured by the Codon Adaptation Index (CAI), suggesting sub-optimal codon frequency does not trigger degradation of Tst-regulated RNAs (Figure S4A)^57,58^. Instead, we found an enrichment of repeating, interspaced cytidine residues in the coding sequence (CDS), but not in the 5’UTRs or 3’UTRs, of Tst-regulated RNAs (Figure 4A), suggesting cytidine tracts might recruit Tst. To investigate this hypothesis, we compared three actin paralogs (*act42A, act57B* and *act87E*), which were upregulated upon the loss of *tst*, to a fourth, *act5C*, which was not upregulated. The *actin* coding sequences are highly similar (>84% nucleotide identity) and have similar CAIs (Fig S4B). However, a multiple sequence alignment revealed a repeating cytidine tract in the codon wobble position of *act42A, act57B* and *act87E* that is interrupted by purines in the non Tst-target *act5C* (Figure 4B).

**Figure 4.**
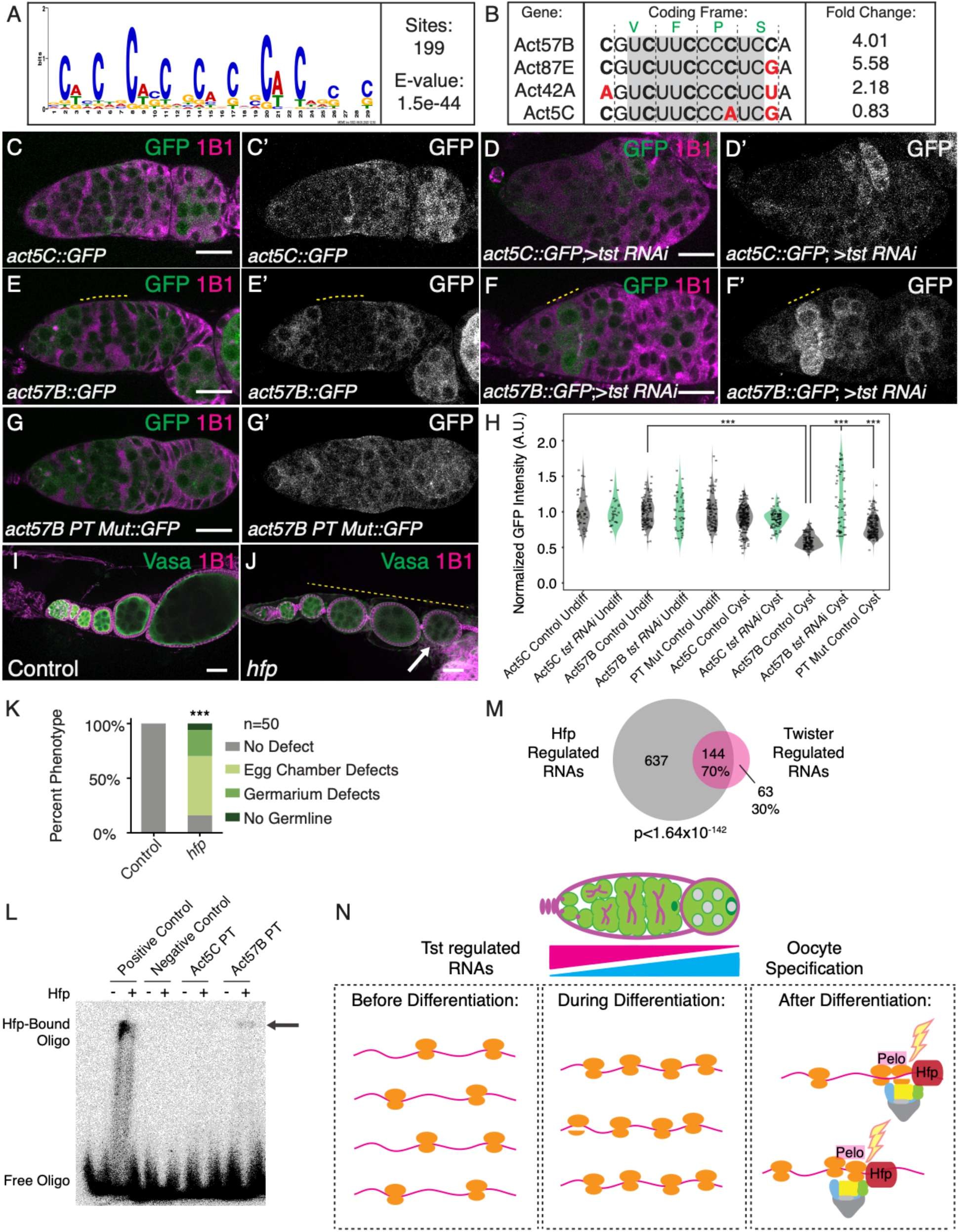
Tst-regulated RNAs are regulated by polypyrimidine rich sequence in their CDS that can be bound by Half Pint. (**A**) MEME logo of the polypyrimidine-rich motif enriched in the CDS of Tst-regulated RNAs. (**B**) CDS alignment of polypyrimidine-rich tracts (PTs) found in the Tst-regulated RNAs *act57B, act87E, act42A*, the non-target paralog *act5C* and their respective fold changes upon loss of *tst*. Black vertical lines indicate coding frame with amino acid symbols above. Probe for EMSA experiments boxed in gray and purine transitions in *act5C* highlighted in red. (**C-C’**) Germarium showing the expression of germline *actin5C::GFP* fusion reporter (green, grayscale) in control and (**D-D’**) *tst RNAi* stained with 1B1 (magenta) that does not change in the cysts. (**E-E’**) Germarium showing the expression of germ line *actin57B::GFP* fusion reporter (green, grayscale) and stained with 1B1 (magenta) showing a decrease in *actin57B::GFP* expression level in cysts (yellow dashed line) and in (**F-F’**) *tst RNAi* showing higher *actin57B::GFP* expression level in cysts (yellow dashed line) compared to control. (**G-G’**) Germarium showing the expression of germ line *actin57B PT Mutant::GFP* fusion reporter (green, grayscale) in WT stained with 1B1 (magenta) showing consistent *actin57B PT Mutant::GFP* in both undifferentiated and cyst stages. (**H**) A.U. quantification of reporter GFP intensity in undifferentiated cells and cyst stages in WT indicating significantly lower *actin57B::GFP* expression in WT cysts compared to undifferentiated cells. Expression of *actin57B::GFP* is significantly higher in *tst RNAi* cysts compared to WT control cysts. Expression of *actin57B PT Mutant::GFP* is significantly higher than *actin57B::GFP* in WT control cysts. (*actin57B::GFP* Control undifferentiated cells n=110, *actin57B::GFP* Control Cyst n=116, p<0.0001, *actin57B::GFP tst RNAi* Cyst n=53, p<0.0001, *actin57B PT Mutant::GFP* Cyst n=158, p<0.0001 Student’s t-Test). (**I**) Confocal image of Control and (**J**) *hfp* mutant ovariole stained with 1B1 (magenta) and Vasa (green) showing egg chambers that do not grow in size (yellow dashed line) and dying egg chamber (arrow). (**K**) Quantification of *hfp* oogenesis defect phenotypes compared to control (n=50, p<0.001, Chi Square Analysis). (**L**) EMSA of recombinant Hfp N-terminal RRMs shows that Hfp RRMs bind the *Drosophila* consensus polypyrimidine-rich sequence (Positive Control) and *act57B* PT sequence (from **B**) but not the random scramble sequence (Negative Control), or the *act5C* PT sequence. (**M**) Venn diagram illustrating overlap of 70% of Tst-regulated RNAs upregulated upon loss of *hfp* (p<1.64×10^−142^, Hypergeometric Test). Scale bars are 10μm. (**N**) In undifferentiated cells Tst-regulated RNAs are highly expressed, yet lowly associated with ribosomes, and required for early oogenesis. During differentiation, translation of Tst-regulated RNAs increases. After differentiation, during oocyte specification, Hfp protein binds in the CDS of Tst-regulated RNAs leading to targeting by Pelo, and the Ski complex.

To evaluate the effect of these sequence differences *in vivo*, we built reporters with the GFP open reading frame fused to the CDS of non-target *act5C* and target *act57B*, as well as a version of *act57B* with cytidine tracts mutated to match *act5C* (PT-mutant). We expressed these reporters under the control of the maternal germline promoter *pgc*, as well as the 3’UTR of *nos*, and 5’UTR of *K10* (Figure S4C-C’’), which are not translationally repressed^32,59-61^. Levels of *act5C-GFP* did not significantly change during oocyte specification or upon loss of *tst* (Figure 4C-D’, H). In contrast, the levels of *act57B-GFP* were significantly reduced in WT cysts compared to undifferentiated cells (Figure 4E-E’, H); we note that the reporter was re-expressed in the egg chambers, which could arise from the strong maternal germline promoter or additional layers of control on Tst-regulated RNAs. Strikingly, upon germline depletion of *tst*, the levels of *act57B-GFP* were strongly elevated in cysts (Figure 4E-F’, H). Expression of *act57B PT Mutant-GFP* was significantly higher in cysts compared to that of *act57B-GFP* (Figure 4G-H), matching *act5C-GFP* and demonstrating the importance of the cytidine tract in promoting destabilization of Tst-regulated RNAs during the cyst stages.

To identify the factor that recruits Tst to these cytidine tracts, we looked for polypyrimidine tract binding proteins (PTBs) expressed during oogenesis, and found two: *hephaetus* (*heph*) and *half pint* (*hfp*), the homolog of human PUF60. While loss of *heph* did not phenocopy *pelo* and *tst*, loss of *hfp* partially phenocopied the oogenesis defects of *pelo* and *tst* (Figure 4I-K)^62,63^. Consistent with previous reports, we found that Hfp is present in the nucleus, where it has been shown to regulate splicing and we also observed it in the cytoplasm, suggesting it can affect RNA stability as well as translation (Figure S4D-E)^63^. To determine if Hfp preferentially binds to the cytidine tracts found in Tst-regulated RNAs, we performed Electrophoretic Mobility Shift Assays (EMSA) using a recombinant protein composed of the two N-terminal RNA Recognition Motifs (RRM) of Hfp, which dimerize on a denaturing gel (Figure S4F-G). We observed that the Hfp RRMs bound the *act57B* PT more efficiently than the *act5C* PT sequence (Figure 4L). To determine if Hfp bound to Tst-regulated RNAs *in vivo*, we immunoprecipitated Hfp from young WT ovaries followed by qRT-PCR. We found that the Tst-regulated RNA *act57B* was robustly associated with Hfp, whereas the non-targets *polar granule component* (*pgc*) and *act5C* were not (Figure S4H). Lastly, to determine if Hfp also co-regulates Tst-regulated RNAs, we performed RNA-seq of *hfp* mutant ovaries. We found that 70% of the RNAs upregulated in *tst*-depleted ovaries were also upregulated >2-fold in *hfp* mutants, including *act57B, act42A* and *act87E* (144/207, Figure 4M), whereas *act5C* was not upregulated in either mutant. We did not observe splicing defects of Tst-regulated RNAs in *hfp* mutants, ruling out mis-splicing as the reason for their upregulation (Figure S4I). Taken together, our data suggest that Hfp binding to a subset of Tst-regulated RNAs can elicit their degradation mediated by both Pelo and Tst by presumably modulating ribosome association.

Having elucidated the mechanisms underlying how Tst-regulated RNAs are recognized for degradation, we sought to determine how correct temporal regulation of these RNAs contributes to oogenesis. We first assessed the functions of Tst-regulated RNAs prior to their degradation. Individual depletion of 6 out of 50 Tst-regulated RNAs we tested using germline-specific RNAi resulted in germline defects, including a complete loss of the germline (Supplemental Table 2) (Figure S5A-E). Finally, to elucidate why ectopic persistence of Tst-regulated RNAs interferes with later oogenesis, we examined *tst* mutants for hallmarks of oogenesis defects. We did not find any changes in differentiation or nurse-cell endocycling (Figure S6A-F)^11,17,64-67^. However, we did observe that Egalitarian (Egl), a protein required to transport the maternal RNA contribution to the oocyte, always localized to the oocyte in WT but in *tst, pelo* and *hfp* mutants, while Egl initially localized to the oocyte, this localization was not maintained in later egg chambers (Figure S6G-J’)^13,68^. This suggests that targeted RNA degradation is required for proper inheritance of the maternal contribution, which is necessary for oocyte specification. Thus, some Tst-regulated RNAs play critical roles in germ stem cell maintenance, but are detrimental to the transition to a mature oocyte.

In conclusion, we find that specific RNAs expressed during the undifferentiated stages of oogenesis are degraded during oocyte specification preventing them from being inherited as part of the maternal contribution mediated by the NGD components Pelo and Tst (Figure 4N). Aberrant persistence of these RNAs results in loss of oocyte maintenance and death of egg chambers. This suggests that precise curation of the maternal contribution is tightly coupled to successful egg production. Based on our observations, we propose that a germ cell-to-maternal transition (GMT) occurs during oocyte specification. We speculate that the GMT exists to enable both the transition from germ cell to oocyte identity and the accrual of the maternal RNA contribution to the embryo. After fertilization, the maternal contribution is subsequently cleared during the oocyte-to-embryo and maternal-to-zygotic transitions (MZT) to promote a zygotic identity^69-71^. Thus, RNA degradation bookends an oocyte’s fate, regulating both its initiation and termination.

## Materials and Methods

### Fly lines

Flies were grown at 25°C and dissected between 1-3 days post-eclosion. The following RNAi stocks were used in this study: *tst RNAi* (Bloomington #55647), *CG8777 RNAi* (Ski3, VDRC #v100948), *CG3909 RNAi* (Ski8, VDRC #12758), *bam RNAi* (Bloomington #33631), *UAS-Tkv* (Bloomington #35653), *bam RNAi; hs-bam*^33^. The following tissue-specific drivers were used in this study: *UAS-Dcr2;nosGAL4* (Bloomington #25751), *UAS-Dcr2;nosGAL4;bamGFP, If/CyO;nosGAL4* (Lehmann Lab) *nosGAL4;MKRS/TM6* (Bloomington #4442), and *tjGAL4/CyO* (Lehmann Lab)^23^. The following stocks were used in this study: *y*^*1*^*w*^*1118*^*P{ry[+t7.2]=70FLP}3F* (Bloomington #6420), *w*^*1118*^;*Mi{ET1}tst*^*MB10212*^*/TM6C,Sb1* (Bloomington #29100), *hfp*^*9*^,*cu/TM2* (Schüpbach Lab), *M{UAS-hfp.ORF.3xHA}ZH-86Fb* (FlyORF, F000989), *CG8777*^*MI02824*^*/CyO* (Bloomington #35904), *w[1118];Df(2R)ED1770,P{w[+mW.Scer\FRT.hs3]=3’.RS5+3.3’} ED1770/SM6a* (Bloomington #9157), *w[1118]; Df(3R)Exel9013/TM6B, Tb[1]* (Bloomington #7991), *pelo*^*1*^*/CyO* (Bloomington #11757).

### Genotypes used to enrich specific stages of germline

Germline Stem Cells: *nosGAL4>UAS-Tkv* (Bloomington #35653)^9,32,72^. Cystoblasts: *nosGAL4>bam RNAi* (Bloomington #33631)^11,64,65^. Differentiating Cysts: *nosGAL4>bam RNAi; hs-bam*^33^. Female flies were heat shocked at 37° C for 2 hours, incubated at room temperature for 4 hours and heat shocked again for 2 hours. This was subsequently repeated the next day and flies were dissected. Young Wild Type: *y*^*1*^*w*^*1118*^*P{ry[+t7.2]=70FLP}3F* (Bloomington #6420). Female flies were collected and dissected within 2 hours of eclosion.

### Dissection and Immunostaining

Flies were dissected in 1X PBS and samples were fixed for 10 minutes in 5% methanol-free formaldehyde^32^. Ovary samples were washed in 1 mL PBT (1X PBS, 0.5% Triton X-100, 0.3% BSA) 4 times for 7 minutes each. Primary antibodies were added in PBT and incubated at 4°C rotating overnight. Samples were washed 4 times for 7 minutes each in 1 mL PBT, and once in 1 mL PBT with 2% donkey serum (Sigma) for 15 minutes. Secondary antibodies were added in PBT with 4% donkey serum and incubated at room temperature for 2 hours. Samples were washed 4 times for 7 minutes each in 1 mL of 1X PBST (0.2% Tween 20 in 1x PBS) and incubated in Vectashield with DAPI (Vector Laboratories) for 30 minutes before mounting. The following primary antibodies were used: Mouse anti-1B1 (1:20, DSHB), Rabbit anti-Vasa (1:1000, Rangan Lab), Chicken anti-Vasa (1:1000), Rabbit anti-GFP (1:2000, Abcam, ab6556), Rabbit anti-Blanks (1:1000, Sontheimer Lab), Mouse anti-Actin C4 (Sigma, MAB1501), Rabbit anti-Cleaved Caspase3 (1:300, Cell Signaling #96615), Rabbit anti-Egl (1:1000, Lehmann Lab), Alexa 488-Conjugated Phalloidin (Cell Signaling #8878), Mouse anti-Hfp (1:25, Schüpbach Lab)^35,37,73,74^. The following secondary antibodies were used: Alexa 488 (Molecular Probes), Cy3, and Cy5 (Jackson Labs) were used at a dilution of 1:500.

### Fluorescence Imaging

The ovary tissue samples were visualized under 10X dry, 20X dry and 40X oil objective lenses and images were acquired using a Zeiss LSM-710 confocal microscope. Confocal images were processed with ImageJ. A.U. The images were quantified using ImageJ with the Measurement function.

### Generation of Transgenic Flies

The pCasper2 plasmid containing the *pgc* promoter, *nos* 5’UTR, eGFP CDS and K10 3’UTR was used as a backbone to generate Actin-GFP reporter constructs^32^. gBlocks (IDT) of the *actin5C, actin57B* and *actin57B* PT-Mutant CDSs were individually cloned upstream of GFP by digesting with SpeI (NEB, R0133S). Constructs were ligated through Gibson Assembly (NEB, E2611S), utilizing complementary overhangs between the CDS fragment and the pCasper2 backbone. Injection of these plasmids into *Drosophila* embryos was conducted by BestGene Inc.

### Gateway Cloning

The coding sequence of Tst was PCR amplified from cDNA to include flanking attB sites. BP recombination was carried out according to the manufacturer’s protocol using equimolar amounts (100 fmol) of the attB-PCR product and the pDONR entry clone plasmid (Invitrogen, 12535-019). Components were incubated in TE buffer with BP Clonase enzyme mix and reaction buffer at 25°C for one hour. 2 μg/μL Proteinase K was added to the reaction and incubated at 37°C for one hour. Plasmid was then transformed into DH5α competent cells and plated on LB-Kan plates at 37°C overnight (Thermo, 18265017). Cells of individual colony samples were propagated and plasmid was purified. LR recombination reaction was performed with the pPPW and pPGW destination vectors (Gateway Collection). Components were incubated in TE buffer with LR Clonase enzyme mix and reaction buffer at 25°C for one hour. 2 μg/μL Proteinase K was added to the reaction and incubated at 37°C for one hour. Plasmid was then transformed into XL10-Gold competent cells and plated on LB-Kan plates at 37°C overnight (Integrated Sciences, 200315). Cells of individual colony samples were propagated, plasmid was purified and sequenced to verify insertion.

### RNA Isolation

Ovaries were dissected in 1X PBS and homogenized in 50uL of TRIzol (Invitrogen, 15596026)^32^. RNA was isolated by adding an additional 950 uL of TRIzol and 230uL of Chloroform with mixing. Samples were centrifuged at 13,000 rpm, 4°C for 15 minutes. Aqueous phase was transferred to a new tube, nucleic acids were precipitated using 1 mL of 100% ethanol, 52 μL of 3M Sodium Acetate and precipitated for >1 hour at −20°C. Samples were centrifuged at 13,000 rpm, 4°C for 20 minutes. Ethanol was decanted, pellet was washed with 70% ethanol and dried at room temperature for 10 minutes. Pellet was dissolved in 20 μL RNase free water and placed in a 42°C water bath for 10 minutes. Concentration of nucleic acid samples were measured on a spectrophotometer and treated with DNase (TURBO DNA-free Kit, Life Technologies, AM1907).

### Quantitative Real Time-PCR (qRT-PCR)

1 μL of cDNA was amplified using 5μL of SYBR green Master Mix, 0.3 μL of 10μM of each reverse and forward primers in a 10 μL reaction^32^. The thermal cycling conditions consisted of 50°C for 2 minutes, 95°C for 10 minutes, 40 cycles at 95°C for 15 seconds, and 60°C for 60 seconds. The experiments were carried out in technical triplicate and minimum 2 biological replicates for each sample. *rp49* gene was utilized as a control. To calculate fold change in mRNA levels to *rp49* mRNA levels, average of the 2^ΔCt for the biological replicates was calculated. Error bars were plotted using standard error of the ratios. P-value was determined by Students t-test.

### RNAseq library preparation

Total RNA samples were run on a 1% agarose gel to assess sample integrity^75^. To generate mRNA-Seq libraries, total RNA was incubated with poly(A) selection beads. mRNA enriched sequencing libraries were made with the NEXTflex Rapid Directional RNAseq Kit (BioO Scientific Corp.) and corresponding protocol. mRNA was fragmented at 95°C for 13 minutes to achieve ∼300 bp fragments. 75 bp single-end (or paired-end as specified) mRNA sequencing was performed each sample with an Illumina NextSeq500, carried out by the Center for Functional Genomics (CFG).

### RNA-seq analysis

Sequenced reads were aligned to the *D. melanogaster* genome (UCSCdm6 and FlyBase R6.01) using HISAT2 v2.0.5^76^. Unambiguously mapping reads RefSeq annotated mRNA and lincRNA were quantified using featureCounts v1.5.1 default parameters^77^. Genes with >=0.5 reads per million (RPM) in one of WT ovaries, *tst* mutant ovaries, *tst RNAi* ovaries, or young ovaries were retained for further analysis (N=9251 genes). Tst regulated mRNAs were classified as genes whose transcript-per-million (TPM) expression levels were >4-fold increased in the *tst RNAi* samples; a subset of these that are additionally >2-fold increased in the *tst* mutant samples were considered to be strong Tst regulated mRNAs (N=207 genes). To curate a set of non-target genes to serve as a background set, we identified genes that differed <1.25 fold between WT and *tst RNAi* and selected the 3 non-targets with the most similar *tst RNAi* expression level to each Tst regulated mRNA, to yield a set of 621 non-targets. For polysome profiling samples, normalized ribosome occupancy was calculated as log_2_(polysome TPM / input TPM). To compare ribosome occupancy of Tst regulated mRNAs across different conditions relative to the global differences observed, we normalized Tst regulated mRNA ribosome occupancy by mean non-target occupancy, i.e. log_2_((polysome_target TPM) / (input_target TPM) / average (polysome_non-target TPM / input_non-target TPM)). Codon optimality index (CAI) was calculated for each gene relative to the codon frequencies in the top 100 expressed genes in WT ovaries according to Sharp & Li, 1987^57^.

### EdU

Ovaries were dissected into Schneider’s Media and incubated in 10μM EdU solution (Click-iT EdU Flow Cytometry Assay Kit) rotating for one hour^78^. Samples were fixed in 3.7% formaldehyde in PBS, rotating for 30 minutes. Fixative was then aspirated and samples were washed with 1 mL PBS for 10 minutes and permeabilized in 1 mL of permeabilization solution (1% Triton X-100 in PBST) rotating for 20 minutes. Samples were then washed in 1 mL PBS rotating for 10 minutes. Click-iT reaction cocktail (PBS, CuSO_4_, Fluorescent dye azide and Reaction Buffer Additive) was made according to manufacturer’s directions and added to each sample. Tubes were protected from light and rotated at room temperature for 30 minutes. Samples were then washed once with 1 mL of Click-iT reaction rinse buffer and once with 1 mL PBS. Ovary samples were then transitioned to the immunostaining protocol.

### Fluorescent *in situ* Hybridization

Ovaries were dissected in RNase free 1X PBS and fixed in 1 mL of 5% formaldehyde rotating for 10 minutes. Samples were then washed three times for five minutes each in PT buffer (PBS, 0.1% Triton X-100) and dehydrated in successive methanol washes for six minutes each (30%, 50%, 70%). A final 100% methanol wash was carried out for 12 minutes. Samples were equilibrated to PT buffer by conducting successive methanol washes for six minutes each (70%, 50%, 30%), followed by three PT washes of six minutes each. Ovaries were pre-hybridized for six minutes in 1 mL Wash buffer (10% Deionized Formamide, 2X SSC in RNase Free H_2_O). Alexa-488 fluorescent probe against *pgc* was generated by Stellaris. Hybridization of probes was conducted at 32°C, covered for >16 hours. Samples were then washed six times in Wash buffer for 2 minutes per wash. Samples were then washed twice in 1 mL Wash buffer for 30 minutes at 30°C. Wash buffer was aspirated and incubated in Vectashield for 30 minutes before mounting. *in situ* experiments were repeated more than three times for control and experimental ovaries.

### RNAScope™ Assay

We utilized a modified RNAscope procedure for *Drosophila* ovaries described previously^79^. Probes were designed and generated by Advanced Cell Diagnostics with specificity to target base pairs 29-1250 of *blanks* mRNA (accession number from NCBI: NM_139709.2), base pairs 1196-1693 of *actin57B* mRNA (NM_079076.4). Ovaries were dissected in RNase free 1X PBS and fixed in 1 mL of 5% formaldehyde rotating for 10 minutes. Samples were then washed three times for five minutes each in PT buffer (PBS, 0.1% Triton X-100) and dehydrated in successive methanol washes for six minutes each (30%, 50%, 70%). A final 100% methanol wash was carried out for 12 minutes. Samples were equilibrated to PT buffer by conducting successive methanol washes for six minutes each (70%, 50%, 30%), followed by three PT washes of six minutes each. Ovaries were pre-hybridized for six minutes in 1 mL of RNAScope Wash buffer (ACD, 310091). Hybridization of probes was conducted at 40°C, covered for >16 hours. Samples were then washed three times in RNAscope wash buffer for 5 minutes per wash, fixed in 4% formaldehyde in 1X PBS at room temperature for 10 minutes and washed in buffer three times for 5 minutes each. Ovaries were incubated in a successive series of amplifier solutions (Amp). Amp 1 for at least 45 minutes at 40°C, Amp 2 for 45 minutes at 40°C, Amp 3 for 45 minutes at 40°C, Amp 4 for 45 minutes at 40°C. After each Amp step ovaries were washed in wash buffer 5 times for 3 minutes each at room temperature. Samples were then washed in 1 mL PBT for 5 minutes and mounted in Vectashield. RNAscope experiments were repeated more than three times for control and *tst RNAi* ovaries.

### Materials and reagents

Fly food was made according to previously published procedures, and filled narrow vials (Fisherbrand Drosophila Vials; Fischer Scientific) to approximately 12mL^32^.

### RNA Immuno-Precipitation (RIP)

65 pairs of ovaries were dissected in 1X PBS^32^. After dissection, PBS was aspirated and 100 μl of RIPA buffer (50 mM Tris pH 8.0, 1% Triton X-100, 0.1% sodium deoxycholate, 0.1% SDS, 140 mM NaCl, 1mM EDTA, 1 mM PMSF, 1 protease inhibitor pill per 50 ml) was added and the sample was homogenized. An additional 200 μl of RIPA buffer was added to the lysate and mixed. The lysate was then centrifuged at 13,000 rpm for 10 minutes at 4°C. The supernatant was transferred to a new tube. 10% of the cleared homogenate was set aside as input, 4X SDS buffer was added the sample was heated at 95°C for 5 minutes and stored at −20°C until Western analysis. An additional 10% of homogenate was used for RNA input, 100 μl of TriZol was added, mixed and this sample was stored in −80°C. 40% of homogenate was used for the IgG control and the remaining 40% was used for RIP. The following antibodies were added to the lysate and incubated at 4°C for 3 hours; 1 μl of Rabbit anti-HA (abcam, 9110), 1 μl of ChromePure Rabbit IgG (Jackson ImmunoResearch Labs). Protein A Dynabeads (Thermo Fisher Scientific) were separated into 15 μl aliquots for each sample and washed four times in 400 μl of 1:10 diluted protease inhibitor-containing Net2 buffer (50 mM Tris-Cl [pH 8.0], 150 mM NaCl, 10% NP-40) on a magnetic rack. The beads were then re-suspended in 100 μl of Net2 buffer. After lysate incubation 25 μl of washed beads was added to each sample and incubated overnight at 4°C. Beads were washed six times with 500 μl of 1:10 diluted Net2 buffer for 2 minutes each. Beads were then resuspended in 25 μl of Net2 buffer. An aliquot of 10 μl was used for Western Blot analysis. The remaining 15 μl was used for RNA extraction.

### Protein Purification

The coding sequence of two adjacent RRMs of Hfp (Amino acids 110-326) was PCR amplified from cDNA to incorporate Nco1 and Kpn1 sites and ligated into the pETM-82 expression plasmid via Gibson assembly (NEB)^32^. The completed plasmid was transformed into Rosetta BL21 cells (Millipore Sigma, 70954-3). A starter culture of 5 ml of Rosetta BL21 cells containing the completed plasmid were grown overnight at 37°C in LB with Kanamycin. This culture was then added to 1000 mL of LB-Kan media. Cells were shaken at 220 rpm at 37°C for 3 hours until OD600 ∼0.6. To induce protein expression, 0.5 mM IPTG was added to the culture and then shaken at 220 rpm at 37°C for 3 hours. The cells were then centrifuged at 4000xg for 20 minutes at 4°C in 50 mL aliquots. The pellet was re-suspended in 3 mL of re-suspension buffer (20 mM Na phosphate, 50 mM NaCl, 20 mM imidazole, 10 ul of 500 mg/ml pH 7.4), sonicated on ice at 20% intensity for 20 seconds for 3 pulses using 1/8-inch probe. The suspension was then centrifuged at 10,000xg for 10 minutes at 4°C. The column (His GraviTrap, GE Cat#11-0033-99) was equilibrated with 10 mL binding buffer (20 mM Na phosphate, 50 mM NaCl, 20 mM imidazole, 10 ul of 500 mg/mL pH 7.4). The supernatant was added to the column and washed with increments of 1 mL, 4 mL and 5 mL of binding buffer. The protein was then eluted using the following washes; twice with 1 mL of elution buffer #1, twice with 1 mL of elution buffer #2 and three times with 1 mL of elution buffer #3. Elution Buffer #1: 20 mM NaPO4, 50 mM NaCl, 150 mM imidazole, pH 7.4 Elution Buffer #2: 20 mM NaPO4, 50 mM NaCl, 300 mM imidazole, pH 7.4 Elution Buffer #3: 20 mM NaPO4, 50 mM NaCl, 500 mM imidazole, pH 7.4 The first elution contained purified Hfp RRM protein. The eluted protein was de-salted using the PD-10 column (GE #17-0851-01). The column was equilibrated with 25mL of binding buffer (10 mM HEPES, 150 mM KCl, pH 7.5) and centrifuged at 3500 rpm for 2 minutes. The eluted protein was slowly added to the column and centrifuged at 3500 rpm for 2 minutes. Desalted protein concentration was determined by Bradford assay. The eluted protein was then stored in 20% glycerol at −80°C until use.

### Electrophoretic Mobility Shift Assay (EMSA)

Positive control oligo: 5’-UUUUUCUCUU-3’, negative control scramble: 5’-UACGUACGUA-3’, *act5C* sequence: 5’-UCUUCCCCAUC-3’, *act57B* sequence: 5’-UCUUCCCCUC-3’ RNA oligonucleotides were end-labeled using T4 Kinase (NEB) with ATP [γ-32P]^32^. Excess ATP was removed through G-25 Sephadix Columns (Roche, 11273990001). RNA-binding reactions were performed in 1X Binding Buffer (50mM Tris pH 7.5, 150mM NaCl, 2mM DTT, 0.1mg/μl BSA, 0.001% Tween-20, 0.5μl of dIdC, 1μl RNaseOUT and 0.5μl of yeast t-RNA)^80^. 3.0 nM of RNA oligo and 3.6μM purified Hfp RRM protein was incubated for 20 minutes at RT and then ran on an 3.5% native polyacrylamide TBE gel at 150V for 4 hours at 4°C. The gel was then dried onto Whatmann filter paper for one hour and exposed to a phosphor screen overnight. A Typhoon Trio imager was used to image the phosphor screen.

### Subcellular Fractionation

50 adult Wild Type ovaries were dissected in 1X PBS and homogenized with 10-20 strokes of a plastic homogenizer in 100 μL hypotonic lysis buffer (10mM HEPES pH 7.9, 1.5 mM MgCl_2_, 10 mM KCl, 0.5 mM DTT). Homogenate was incubated on ice for 15 minutes. 50 μL of homogenate was aliquoted in a new tube, 4X SDS buffer was added, sample was boiled at 95°C for 5 minutes and stored in −20°C until use as total homogenate. The remaining homogenate was centrifuged for 10 minutes at 1000g. 50 μL of supernatant was collected 4X SDS buffer was added, sample was boiled at 95°C for 5 minutes and stored in −20°C until use as cytoplasmic fraction. The pellet was resuspended in high salt extraction buffer (20mM HEPES pH 7.9, 25% glycerol, 420 mM NaCl, 1.5 mM MgCl_2_, 0.2 mM EDTA, 0.5 mM DTT) and centrifuged for 5 minutes at 20,000g. Supernatant was collected 4X SDS buffer was added, sample was boiled at 95°C for 5 minutes and stored in −20°C until use as total nuclear fraction.

### Western Blot

Twenty wild-type size ovaries or 40 mutant size ovaries were dissected in 1X PBS^32^. After dissection, PBS was aspirated and 30 μl of NP-40 buffer with protease inhibitors added to the tissue and homogenized. The lysate was centrifuged at 13,000 rpm for 15 minutes at 4°C. Aqueous layer was transferred into a new tube while avoiding the top lipid layer. 1 μl of the protein extract was used to carry out a Bradford (Bio-Rad, 500-0205) assay. 25 μg of protein was denatured with 4X Laemmli Sample Buffer (Bio-Rad, 161-0747) and β-marcepthanol at 95°C for 5 minutes. The samples were loaded in a Mini-PROTEAN TGX 4-20% gradient SDS-PAGE gels (Bio-Rad, 456-1094) and run at 110V for 1 hour. The proteins were then transferred to a 0.20 μm nitrocellulose membrane at 100V for 1 hour at 4°C. After transfer, the membrane was blocked in 5% milk in PBST for 2 hours at RT. The following antibodies were used: Mouse anti-Hfp (1:1000, Schüpbach Lab), Rabbit anti-Orb (1:1000, Lehmann Lab), Rabbit anti-His (1:000, Rockland Inc., 600-401-382), Mouse anti-Fibrillarin (1:25, DSHB). Primary antibody was prepared in 5% milk in PBST was added to the membrane and incubated at 4°C overnight. The membrane was then washed three times in 0.5% milk PBST. Anti-Rabbit HRP (1:10,000, Abcam, ab97046) or Anti-Mouse HRP (1:10,000, Abcam, ab6721) was prepared in 5% milk in PBST, and was added to the membrane and incubated at room temperature for 2 hours. The membrane was then washed 3 times in PBST. Bio-rad chemiluminescence ECL kit (1705061) was used to image the membrane.

### Egg Laying Assay

Newly eclosed flies were collected and fattened overnight on yeast. Assays were conducted in cages on apple juice plates containing 6 control or experimental females crossed to 4 Wild Type control males. Cages were maintained at 25°C and plates changed daily for counting. Analyses were performed on three consecutive days. Total number of eggs laid was counted and averaged. Both control and experimental experiments were conducted in triplicate.

### Polysome profiling and Polysome-Seq

30 Wild Type or 150 experimental ovary pairs were dissected in 1X PBS and immediately flash frozen on liquid nitrogen^32,81^. Samples were homogenized in lysis buffer and 20% of lysate was used as input for mRNA isolation and library preparation (as described above). Samples were loaded onto 10-45% CHX supplemented sucrose gradients in 9/16 x 3.5 PA tubes (Beckman Coulter, #331372) and spun at 35,000 x g in SW41 for 3 hours at 4°C. Gradients were fractionated with a Density Gradient Fractionation System (#621140007). RNA was extracted using acid phenol-chloroform and precipitated overnight. Pelleted RNA was resuspended in 20 μL water, treated with TURBO DNase and libraries were prepared as described above.

### MEME Analyses

The 5’UTR, CDS, 3’UTR and full transcript sequences of all 207 Tst-regulated target genes were individually analyzed by the MEME algorithm^82^. Classic mode analysis was utilized to conduct *de novo* motif search with default parameters as well as Any Number of Repetitions (anr) mode. Discriminative mode analysis was conducted against 621 non-target gene sequences as background with default parameters. Motif logos, number of sites, and p-values all reported as produced by output of the program.

## Supporting information

Supplemental dataupplemental Files 1

## Acknowledgements

We would like to thank all members of the Rangan Lab as well as Dr. Sano H, Juliano C, Belfort M, and Farrell J for discussion and comments on the manuscript. We would also like to thank the Schüpbach Lab for the Hfp antibody and mutant flies, the Sontheimer Lab for the Blanks antibody, and the Newbury Lab for flies and reagents. P.R. is funded by the NIH/NIGMS (R01GM111779-06). P.B. is funded by NIH (grant 1F31GM126784-01) and by the RNA Institute. M.T.L. was supported by start-up funds from the Univ. of Pittsburgh.

---

**Figure S1.**
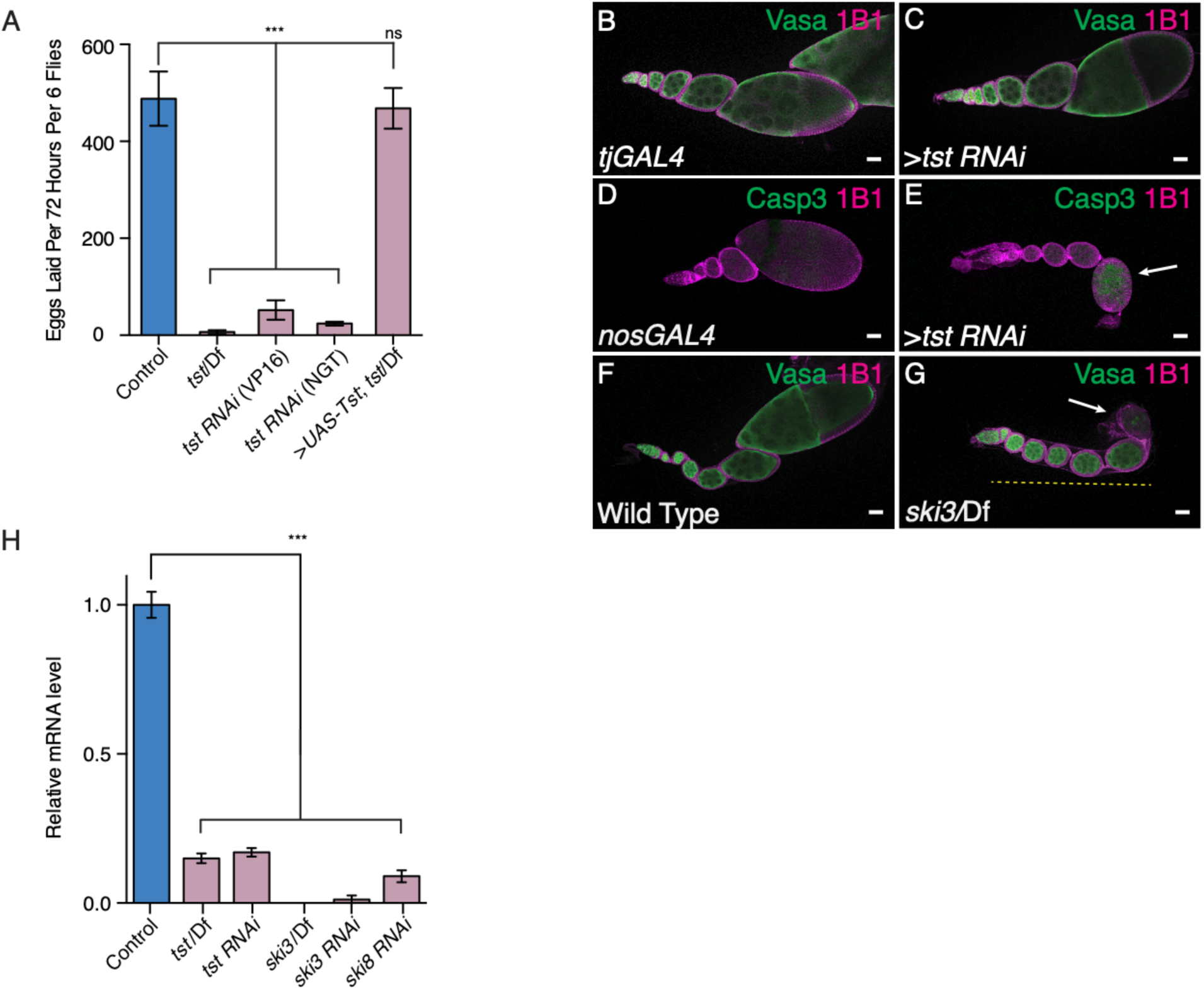
Ski Complex components are required for successful oogenesis. (**A**) Egg laying test assaying the fertility of several Ski complex mutants and germline RNAi knockdown females indicating a loss of fertility compared to control (Control vs *tst*/Df n=3, p<0.001, Control vs *tst RNAi (VP16)*, n=3, p<0.001, Control vs *tst RNAi (NGT)*, n=3, p<0.001, Control vs *UAS-Tst;tst*/Df, n=3, not significant (ns) p>0.05, Error bars are standard deviation (SD), Student’s t-Test). (**B**) *tjGAL4* driver control and (**C**) *tst RNAi* ovarioles stained with Vasa (green) and 1B1 (magenta) exhibiting ovarioles that grow in size and generate later stages. (**D**) *nosGAL4* driver control and (**E**) *tst RNAi* ovarioles stained with cleaved Caspase 3 (green) and 1B1 (magenta) indicating dying egg chamber (arrow). (**F**) WT control and (**G**) *ski3* mutant ovarioles stained with Vasa (green) and 1B1 (magenta) indicating egg chambers that do not grow in size (yellow dashed line) and dying egg chambers (arrow). (**H**) qRT-PCR assaying the levels of *tst, ski3* and *ski8* in their respective mutant background or germline RNAi normalized to control levels and indicating successful knockdown (*tst* Control vs *tst*/Df n=3, p<0.001, *tst* Control vs *tst RNAi* n=3, p<0.001, *ski3* Control vs *ski3*/Df n=3, p<0.001, *ski3* Control vs *ski3 RNAi* n=3, p<0.001, *ski8* Control vs *ski8 RNAi* n=3, p<0.001, Error bars are SEM, Student’s t-Test). Scale bars are 10μm.

**Figure S2.**
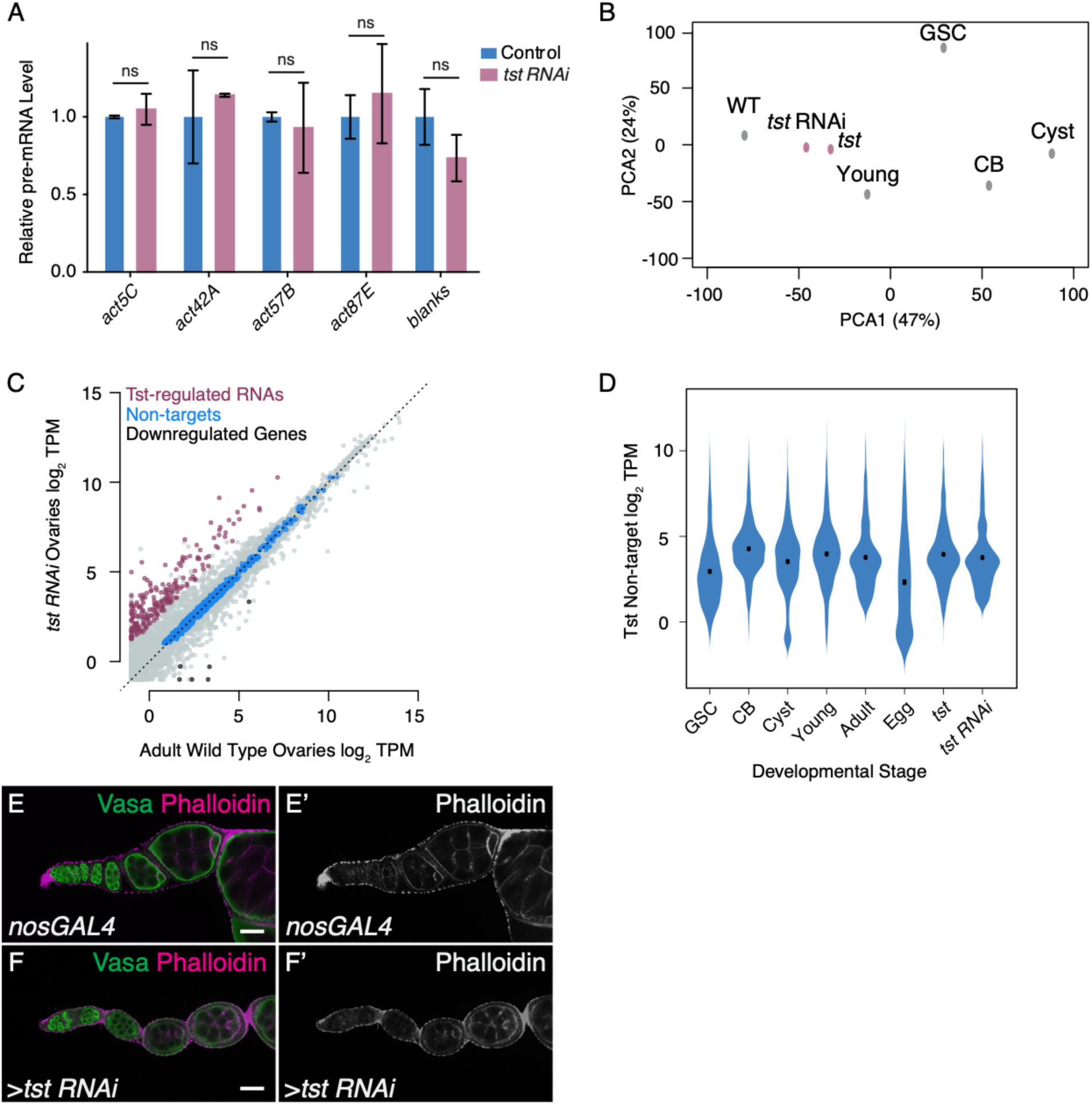
Tst post-transcriptionally regulates its target RNAs. (**A**) qRT-PCR assaying the pre-mRNA levels of several Tst-regulated target genes, including *blanks, act42A, act57B, act87E* and the non-target *act5C* in Control and germline *tst RNAi* normalized to control levels and indicating similar pre-mRNA levels in both conditions (*act5C* pre-mRNA Control level vs *tst RNAi* n=2, ns, p>0.05, *act42A* pre-mRNA Control level vs *tst RNAi* n=2, ns, p>0.05, *act57B* pre-mRNA Control level vs *tst RNAi* n=3, ns, p>0.05, *act87E* pre-mRNA Control level vs *tst RNAi* n=2, ns, p>0.05, *blanks* pre-mRNA Control level vs *tst RNAi* n=3, ns, p>0.05, Error bars are SEM, Student’s t-Test). (**B**) Principal Component Analysis (PCA) comparing several ovary RNA-seq data sets including, adult (WT), *tst RNAi, tst* genomic mutant (*tst*), young WT (Young), germline stem cell enriched (GSC), undifferentiated cystoblast enriched (CB), and differentiating cyst enriched (Cyst). This indicates that the *tst* mutant and *tst RNAi* samples are similar to Adult WT. (**C**) Biplot of RNA-Seq data from Adult WT and *tst* germ line RNAi knockdown ovaries in log_2_ Transcripts Per Million (TPM) highlighting upregulated Tst-regulated RNAs (magenta), non-target RNAs (blue), and RNAs concordantly downregulated in both *tst RNAi* and *tst* genomic mutant ovaries (black). (**D**) Violin plot assaying the expression of non-target genes that do not substantially change in several RNA-seq data sets including Germline Stem Cell enriched (GSC), undifferentiated Cystoblast enriched (CB), and differentiating cyst enriched (Cyst), young WT (Young), adult (WT), unfertilized eggs (Egg), *tst* genomic mutant (*tst*), and germline *tst RNAi* (*tst RNAi*). (**E-E’**) *nosGAL4* driver control and (**F-F’**) *tst RNAi* ovarioles stained with Vasa (green) and Phalloidin (magenta and grayscale) indicating similar levels of phalloidin staining. Scale bars are 10μm.

**Figure S3.**
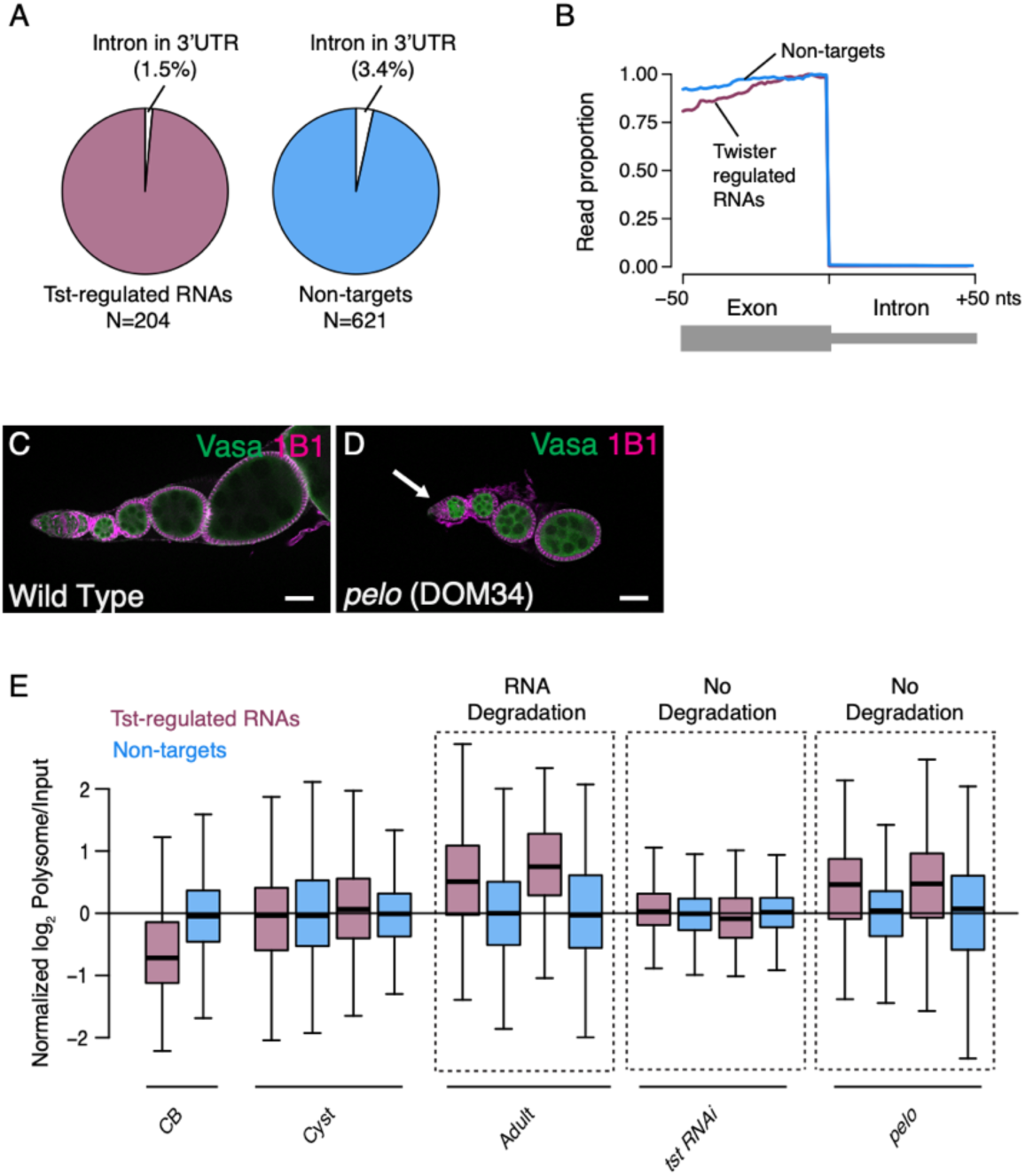
Tst-regulated RNAs exhibit hallmarks of NGD, but not NMD. (**A**) Pie graphs showing the percent of Tst-regulated RNAs (magenta) and non-target mRNAs containing an intron in their 3’UTR indicating that a smaller proportion of Tst-regulated RNAs contain an intron in their 3’UTR (1.5%) compared to non-target RNAs (3.4%). (**B**) Metaplots showing the proportion of RNA-Seq coverage mapping to exon-intron boundaries for both Tst-regulated targets (magenta) and non-targets (blue) indicating that both Tst-regulated RNAs and non-target RNAs are spliced correctly. (**C**) WT control and (**D**) *pelo* mutant ovarioles stained with Vasa (green) and 1B1 (magenta) indicating loss of GSC phenotype (arrow). Scale bars are 10μm. (**E**) Quantification of Normalized log_2_ polysome/input mRNA of Tst-regulated RNAs (magenta), and non-targets (blue) in CB, cyst, adult, *tst RNAi* and *pelo* samples indicating that ribosome association of Tst-regulated RNAs is dynamic during development, but not for non-target RNAs.

**Figure S4.**
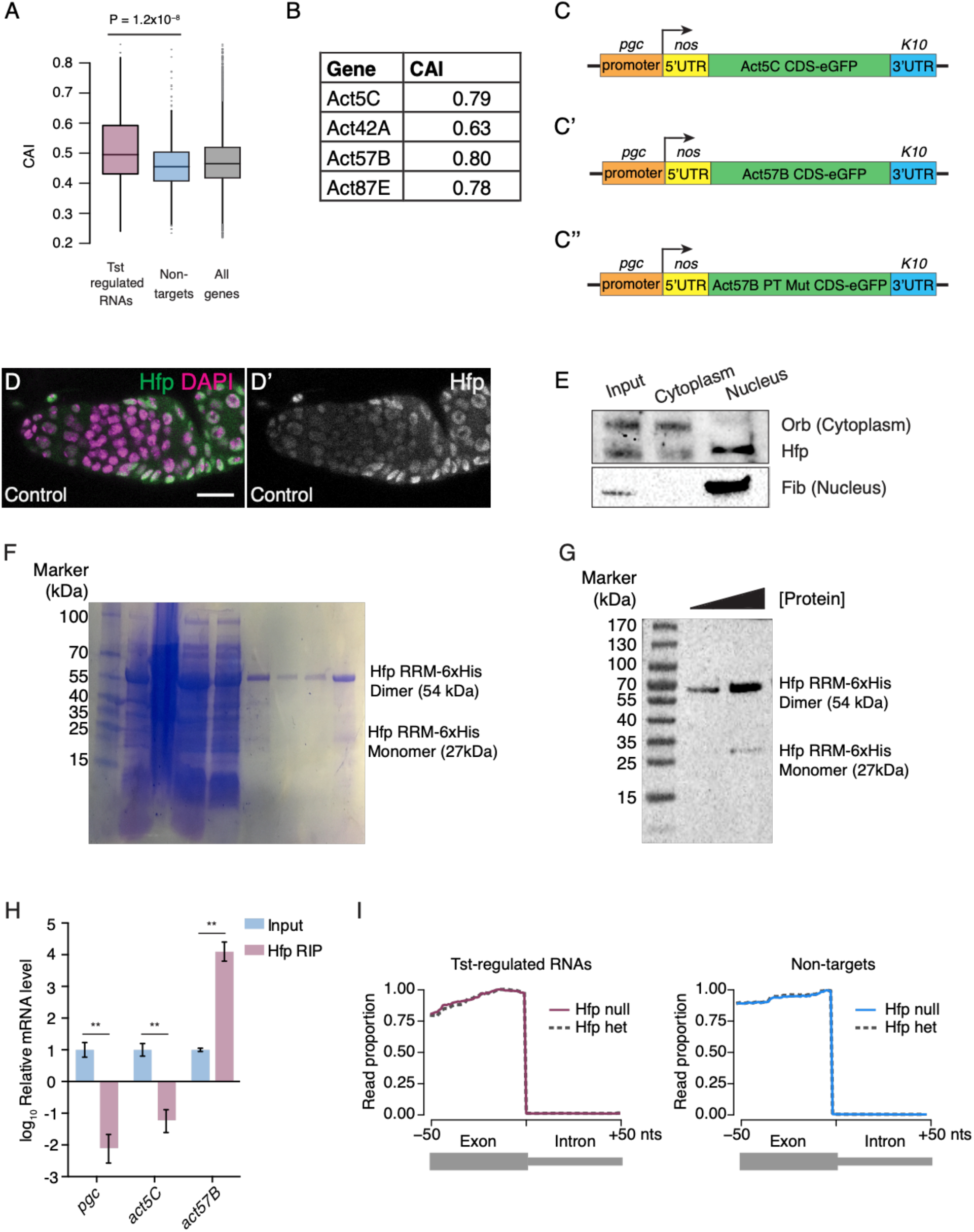
Tst-regulated RNAs are bound by Hfp and do not exhibit suboptimal codon usage. (**A**) Codon Adaptation Index (CAI) comparison for Tst-regulated RNAs (magenta) versus non-targets (blue) indicating a higher CAI for Tst-regulated RNAs (Wilcoxon rank sum test); and all genes (gray). (**B**) Table of the Actin paralog genes and their respective CAI values indicating that they are all very similar. (**C**) Schematic of the *act5C::GFP*, (**C’**) *act57B::GFP* and (**C’’**) *act57B PT Mutant::GFP* reporters under the control of a germline promoter (*pgc*) and 5’UTR (*nos*), and a neutral 3’UTR (*K10*). (**D-D’**) Control germaria stained for Hfp (green and grayscale) and DAPI (magenta) indicating cytoplasmic Hfp expression during early oogenesis. (**E**) Subcellular fractionation Western blot analysis of input, cytoplasm and nucleus for Hfp, Orb and Fibrillarin indicating that Hfp is present in both the nucleus and cytoplasmic fractions. (**F**) SDS-PAGE of a protein marker (lane 1), bacterial supernatant (lane 2), pellet (lane 3), washes (lanes 4-6), and elutions (lanes 7-9) of the Hfp-RRM protein purification process. (**G**) Western blot analysis of the Hfp-RRM 6X-His Tag showing both monomer and dimer bands. (**H**) Hfp-HA RIP and qRT-PCR analyses indicating a de-enrichment of non-target *pgc* and *act5C* levels and an enrichment of target *act57B* levels in Hfp RIP samples compared to input (*pgc* Input vs Hfp-IP n=2, p<0.008, *act5C* Input vs Hfp-IP n=2, p<0.005, *act57B* Input vs Hfp-IP n=2, p<0.005, Error bars are standard error of the mean (SEM), Student’s t-Test). (**I**) Metaplot of the proportion of RNA-seq coverage mapping to exon-intron boundaries in *hfp* mutant and control (heterozygous) RNA-seq data sets for both Tst-regulated RNAs (magenta) and non-targets (blue) indicating correct splicing in both samples. Scale bars are 10μm.

**Figure S5.**
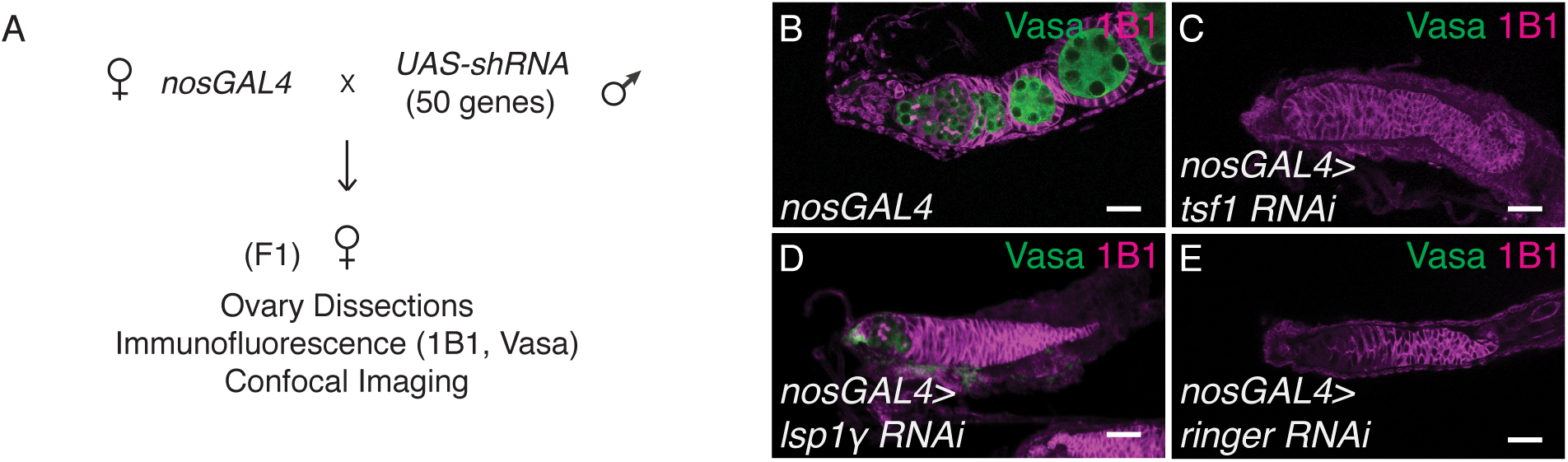
A subset of Tst-regulated RNAs are required for oogenesis. (**A**) Schematic of the germ line RNAi knockdown screen of Tst-regulated genes. 50 Tst-regulated genes were individually depleted by RNAi in the germline by the *UAS-GAL4* system and *nosGAL4* driver. F1 ovaries were dissected and phenotypes were assessed by 1B1 and Vasa staining and confocal imaging. (**B**) *nosGAL4* driver control, (**C**) *tsf1 RNAi*, (**D**) *lsp1γ RNAi*, and (**E**) *ringer RNAi* stained with Vasa (green) and 1B1 (magenta) each exhibiting a complete loss of germ line. Scale bars are 10μm.

**Figure S6.**
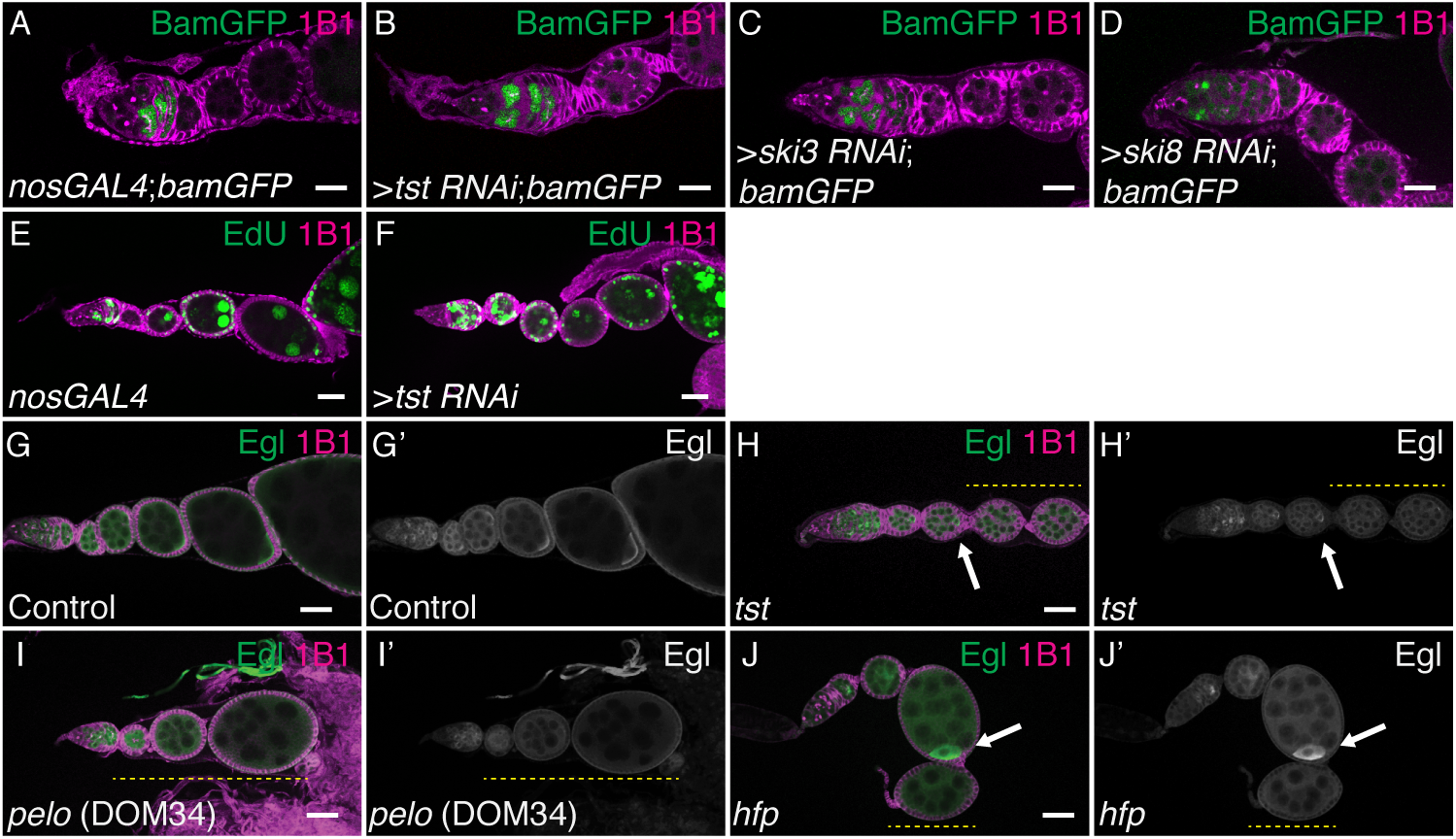
Tst is required for maintaining oocyte fate during oogenesis. (**A**) *nosGAL4;bamGFP* driver control ovariole, (**B**) >*tst RNAi;bamGFP*, (**C**) >*ski3 RNAi;bamGFP*, and >*ski8 RNAi;bamGFP* ovarioles stained with 1B1 (magenta) and GFP (green) indicating appropriate *bamGFP* expression for all samples in the differentiating cells. (**E**) *nosGAL4* driver control and (**F**) >*tst RNAi* ovarioles stained for EdU (green) and 1B1 (magenta) indicating that endocycling is occurring properly. Scale bars are 10μm. (**G-G’**) WT control, and (**H-H’**) *tst RNAi* ovarioles stained with 1B1 (magenta) and Egl (green and grayscale) showing initial localization of Egl (arrow) and subsequent loss of Egl accumulation in *tst RNAi* ovarioles (yellow dashed line). (**I-I’**) *pelo* and (**J-J’**) *hfp* mutant ovarioles stained with 1B1 (magenta) and Egl (green and grayscale) showing initial Egl localization (arrow) and subsequent loss of Egl accumulation (yellow dashed lines).

